# Longitudinal single-cell modeling reveals monocyte reprogramming in juvenile systemic sclerosis following autologous stem cell transplantation

**DOI:** 10.64898/2026.08.21.738279

**Authors:** Julia K. Elrod, Anwesha Sanyal, Theresa Hutchins, F. William Townes, Kathryn S. Torok

## Abstract

Juvenile systemic sclerosis (jSSc) is a rare autoimmune disease marked by skin fibrosis and multi-organ involvement. Autologous stem cell transplantation (ASCT) is an emerging therapy for severe, treatment-refractory jSSc, but its effects on immune cell dynamics remain poorly understood. PBMCs were collected from three patients with jSSc before ASCT and at 6, 12, and 24 months post-ASCT. Patient and healthy control samples were profiled using cellular indexing of transcriptomes and epitopes by sequencing (CITE-seq). We focused on monocytes, given their role in fibrosis-promoting inflammation. To detect longitudinal trends, pseudobulked gene expression (log scale) was regressed against time since ASCT. This approach identified widespread changes in jSSc monocytes, including decreased expression of systemic sclerosis-linked genes, such as *SERPINE1*. On the pathway level, NF-κB-associated inflammatory signaling was elevated in jSSc monocytes at baseline relative to healthy controls and decreased progressively post-ASCT. Genes related to mitochondrial function and oxidative phosphorylation progressively increased in expression after ASCT, suggesting a shift in metabolic state. Compositional changes in monocyte subpopulations were also identified and may have contributed to longitudinal gene expression patterns. Together, these findings characterize the dynamic immune changes in jSSc following ASCT and highlight a widely applicable longitudinal modeling framework for single-cell data.

**Graphical abstract:** 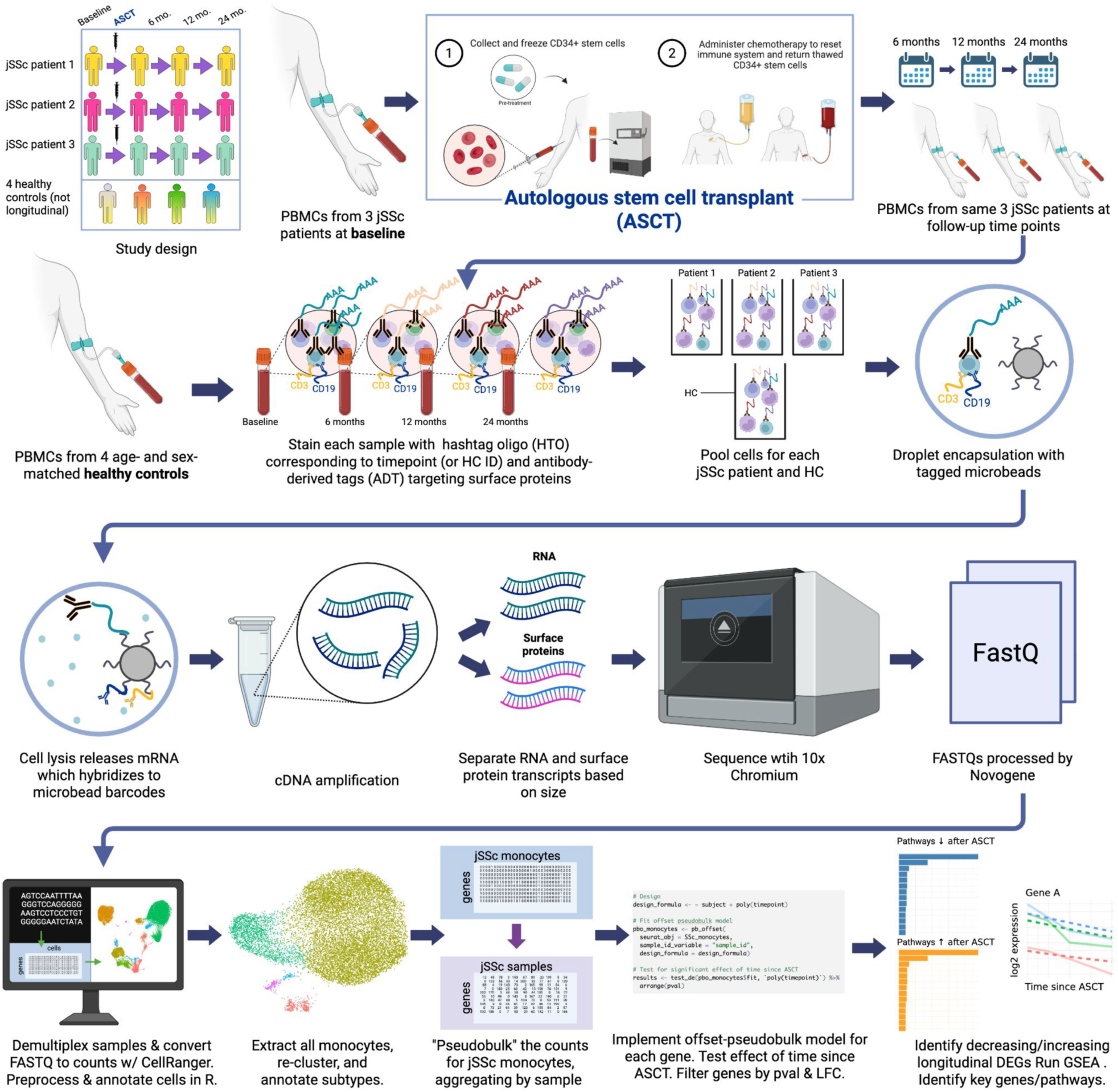

## Introduction

Juvenile systemic sclerosis (jSSc) is a rare and severe autoimmune disease characterized by vasculopathy, immune dysregulation, and progressive fibrosis affecting the skin and internal organs. Although sharing clinical and immunologic features with adult-onset systemic sclerosis, jSSc presents unique challenges related to early disease onset, long-term morbidity, and limited evidence to guide management (1, 2). Early initiation of therapy is recommended, typically including immunosuppressive agents and supportive interventions; however, a subset of patients develop severe, treatment-refractory disease, requiring escalation of care (3).

Autologous stem cell transplantation (ASCT) has emerged as a therapeutic option for severe jSSc with an insufficient response to standard therapies (4). Randomized trials in adults with SSc have demonstrated improved event-free survival and organ-specific outcomes compared with conventional therapy (cyclophosphamide) (5–9). Despite these advances, the application of ASCT in pediatric SSc populations remains limited, and the immunologic mechanisms underlying treatment response, particularly at the level of individual immune cell populations, are incompletely understood. A more detailed characterization of immune changes following ASCT in jSSc is critical to understanding disease pathogenesis and informing future therapeutic strategies.

High-dimensional single-cell approaches offer an opportunity to define immune dysregulation in systemic sclerosis with greater resolution. In particular, circulating monocytes and tissue macrophages have been strongly implicated in fibrotic and inflammatory pathways central to disease pathogenesis (10–15). However, most prior studies rely on cross-sectional data or comparisons between only two time points at once, limiting the ability to characterize dynamic changes in gene expression over time.

Here, we performed longitudinal profiling of peripheral blood mononuclear cells (PBMCs) from patients with jSSc undergoing ASCT using cellular indexing of transcriptomes and epitopes by sequencing (CITE-seq), which integrates transcriptomic and surface protein measurements at single-cell resolution. Focusing on monocytes, we applied a generalized linear model (GLM) framework with a coefficient for time since ASCT to test all genes for statistically significant longitudinal changes, moving beyond the two-way comparisons typical in longitudinal omics studies. This approach adapts the offset-pseudobulk model described by Lee and Han, extending that GLM framework to include a log-linear coefficient for time (16). To our knowledge, this modeling approach has not been commonly applied to longitudinal genomic data. Using this strategy, we characterize concurrent transcriptional and compositional changes across jSSc PBMCs following ASCT, with an emphasis on circulating monocytes, and define a generalizable analytical framework for longitudinal single-cell studies.

## Results

### CITE-seq annotation and cell type proportions in jSSc and healthy PBMCs

PBMCs were profiled by CITE-seq in three patients with jSSc at baseline prior to ASCT and at 6, 12, and 24 months post-ASCT, along with four age- and sex-matched healthy controls. A total of 11 jSSc samples were included in the analysis after exclusion of one sample due to poor quality. The demographics of the three jSSc patients were male White non-Hispanic, female Black non-Hispanic, and female White Hispanic (Supplemental Table 1). Patients were diagnosed between the ages of 7–15 years, and underwent ASCT between the ages of 15–21 years. All exhibited severe disease with significant pulmonary involvement, were refractory to prior therapies, and had a baseline modified Rodnan skin score (mRSS) ranging from 5 to 24 (Supplemental Table 1).

Following quality control and filtering genes with low expression, 28,943 PBMCs and 19,170 genes were retained for analysis (see Methods for further detail). Specific adaptive thresholds used for each library are included in Supplemental Table 2. Multimodal clustering of all patient and healthy control cells, using the Seurat package in R, identified 16 clusters, which were subsequently consolidated into 11 major immune cell types based on canonical RNA and surface protein markers (Figure 1; Supplemental Figure 1; and Supplemental Table 3).

**Figure 1:**
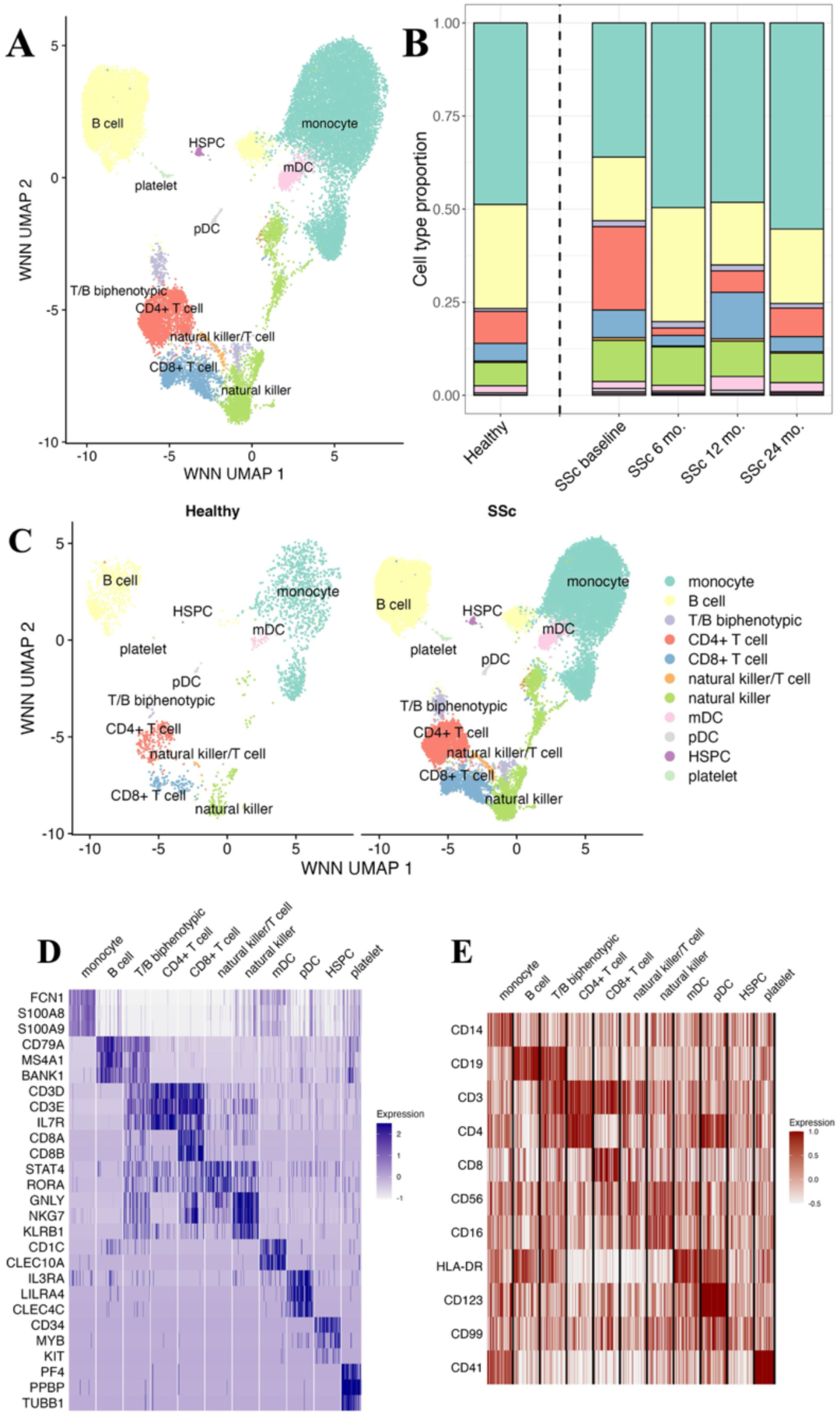
PBMC annotations, cell type proportions, and multimodal markers. A. Multimodal (RNA and surface protein) UMAP of all 28,943 PBMCs from 11 jSSc samples and 4 healthy control samples, annotated with 11 cell types. B. PBMC cell type proportions in healthy controls and in jSSc patients over time since ASCT. Natural killer and T cell populations are expanded in jSSc, and approach healthy control proportions following ASCT. C. UMAPs of all cell types, split into 2,383 healthy cells and 26,560 SSc cells. Compared to SSc cells, very few healthy cells were identified as hematopoietic stem and progenitor cells (HSPC) or T/B biphenotypic cells. D. Heat map showing RNA markers used to identify cell types. E. Heat map showing ADT (surface protein) markers used to identify cell types.

At baseline, jSSc samples demonstrated higher proportions of T cells and natural killer (NK) cells relative to healthy controls (Figure 1B and Supplemental Table 4), consistent with an activated immune phenotype. Over time, following ASCT, cell type distributions showed partial convergence towards those observed in healthy controls. Monocytes comprised a higher proportion of PBMCs than typically expected, a finding which is addressed in the Discussion section.

### Monocyte subtype annotation and longitudinal shifts in monocyte composition

Given their established role in profibrotic inflammation in systemic sclerosis, monocytes were examined in greater detail. A total of 13,906 monocytes were extracted from the PBMCs and re-clustered, yielding 12 sub-clusters that were annotated into five major populations (Figure 2, A– E; Supplemental Figure 2, A–D; and Supplemental Table 5). These included classical CD14+ monocytes, non-classical CD16+ monocytes, and three smaller populations expressing markers of other immune lineages, described below.

**Figure 2:**
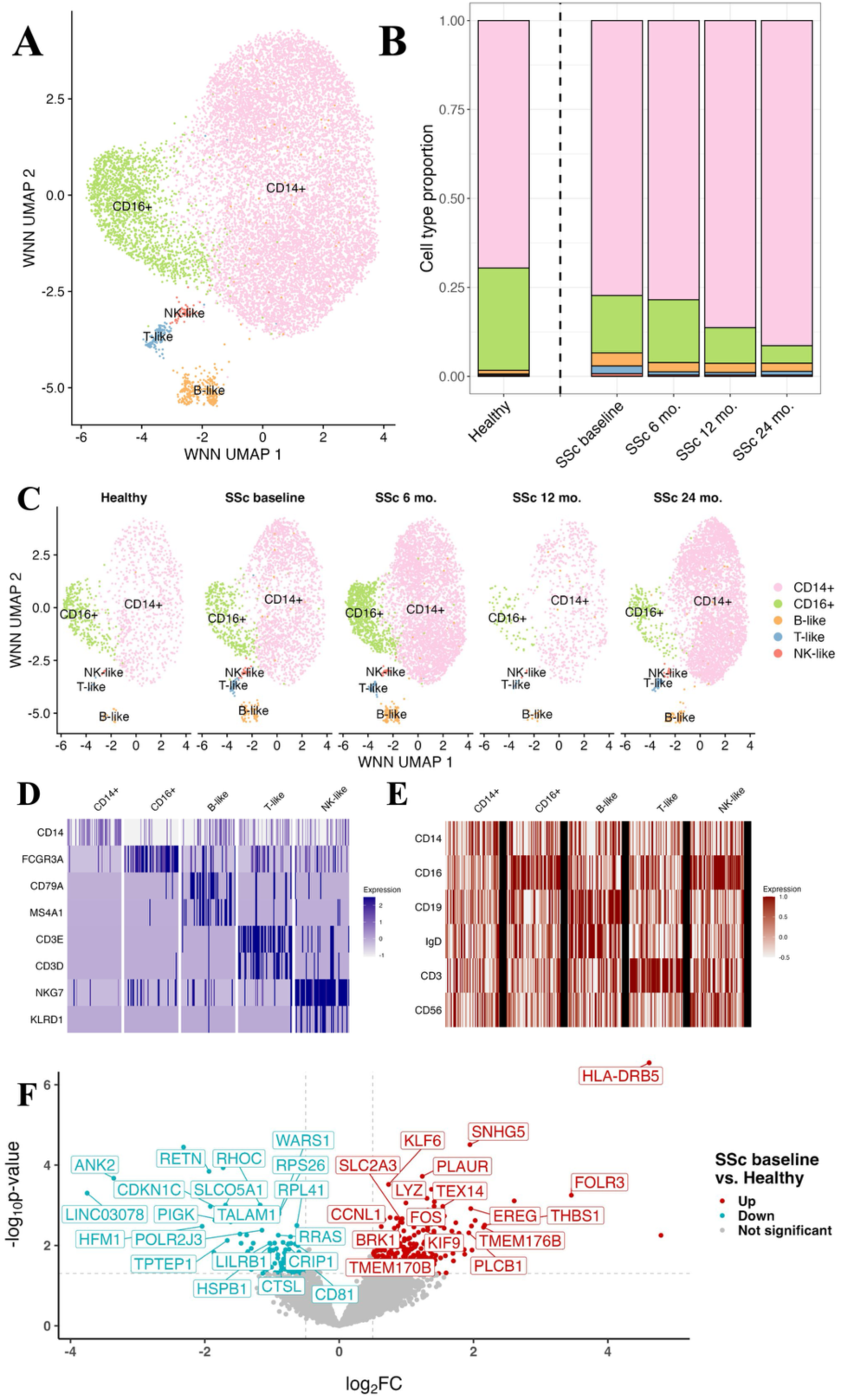
Monocyte subtype annotations, subtype proportions, multimodal markers, and genes with differential expression in jSSc vs. healthy monocytes. A. Multimodal (RNA and surface protein) UMAP of all 13,906 monocytes from 11 jSSc samples and 4 healthy control samples, annotated with 5 subtypes. B. Monocyte subtype proportions in healthy controls and in jSSc patients over time since ASCT. C. UMAPs of monocyte subtypes, split into 1,162 healthy cells, 2,368 cells from SSc patients before ASCT, and 4,892, 819, and 4,665 cells from SSc patients 6, 12, and 24 months after ASCT, respectively. D. Heat map showing RNA markers used to identify monocyte subtypes. E. Heat map showing ADT (surface protein) markers used to identify monocyte subtypes. F. Volcano plot showing genes upregulated or downregulated in jSSc monocytes at baseline compared to healthy controls (*P* < 0.05 and absolute log-fold change (LFC) > 0.5). Genes of biological interest are highlighted.

The relative proportion of classical CD14+ monocytes compared to non-classical CD16+ monocytes was higher in jSSc patients at baseline compared to healthy controls and this difference increased over time following ASCT (Figure 2B and Supplemental Table 6). RNA and protein markers used for annotation are shown in Figure 2, D and E; and Supplemental Figure 2, C–E. Differentially expressed genes in jSSc patients at baseline compared to healthy controls are shown in Figure 2F. Genes of interest that were upregulated in jSSc at baseline compared to healthy controls included *PLAUR*, *THBS1*, *FOS*, and *LYZ*. Equivalent plots comparing jSSc monocytes at 6, 12, and 24 months post-ASCT to healthy controls are included in Supplemental Figure 3, A–C.

### Longitudinal modeling identifies temporal transcriptional changes in jSSc monocytes following ASCT

To characterize temporal changes in monocyte gene expression post-ASCT, we implemented an offset-pseudobulk generalized linear model with a log-linear term for time since ASCT, thus capturing trends in gene expression over time (16). Across jSSc monocytes, hundreds of genes exhibited statistically significant longitudinal changes in expression over the 24-month period following ASCT (Figure 3A and Supplemental Table 7). Notably, several genes implicated in systemic sclerosis pathogenesis, including *SERPINE1*, *FOSL1*, and *SMAD3*, demonstrated some of the strongest negative longitudinal expression trends out of all tested genes (Figure 3B; Supplemental Figure 3D; and Supplemental Table 7) (17–19). Expression trajectories for a wider selection of longitudinal differentially expressed genes (DEGs) are illustrated in Supplemental Figures 4 and 5. Together, these findings indicate broad shifts in transcriptional programs associated with inflammation and fibrosis in circulating monocytes following treatment.

**Figure 3:**
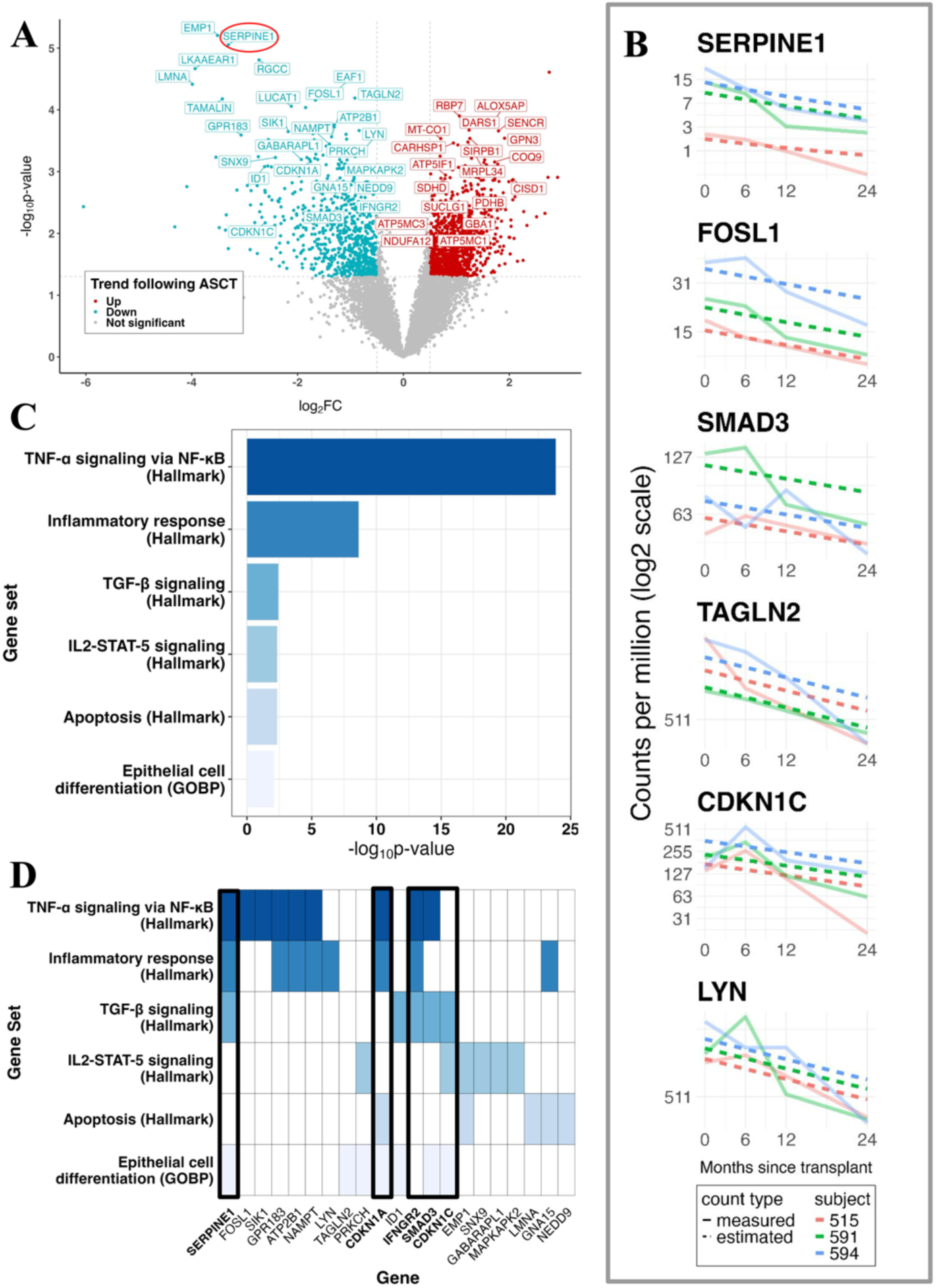
Genes and pathways with longitudinal changes in jSSc monocytes post-ASCT. A. Volcano plot showing genes with increasing or decreasing expression in jSSc monocytes over time since ASCT (*P* < 0.05 and absolute LFC > 0.5). *SERPINE1* (circled) is of special interest due to its suspected role in SSc pathogenesis. B. Longitudinal plots showing gene expression in monocytes over time since ASCT for each patient, as well as fitted expression from our offset-pseudobulk model. C. Gene sets with decreased activity in jSSc monocytes over time since ASCT, selected for biological relevance. D. Top 5 most significantly decreasing genes in each of the pathways in C. Genes appearing in 3+ pathways are highlighted.

### Pathway analysis reveals attenuation of inflammatory and profibrotic signaling

Pathway enrichment analyses of genes with decreasing longitudinal expression identified significant reductions in multiple inflammatory and profibrotic pathways (Figure 3, C and D; and Supplemental Table 8). In particular, nuclear factor-κB (NF-κB)-associated inflammatory signaling and related inflammatory response pathways were elevated in jSSc monocytes at baseline relative to healthy controls and decreased progressively over time following ASCT (Supplemental Figure 6, A and B). These findings suggest a shift towards a less inflammatory transcriptional state and towards that of healthy monocytes over the two years following ASCT.

Additional pathways demonstrating decreased activity longitudinally post-ASCT in monocytes included TGF-β signaling, IL2/STAT5 signaling, apoptosis, and epithelial/mesenchymal-associated cell differentiation programs (Figure 3, C and D; and Supplemental Table 8). To evaluate whether the six main pathways identified (as shown in Figure 3, C and D) represented unique signals, Jaccard similarity was calculated between the set of statistically significant genes associated with each pathway (Supplemental Figure 6C). Limited overlap among most pathway gene sets was identified, indicating largely distinct transcriptional programs, with the exception of partial overlap between the TNF-α signaling via NF-κB gene set and the inflammatory response gene set (Jaccard similarity = 0.17).

In contrast, genes exhibiting increasing expression in jSSc monocytes after ASCT over time were enriched for pathways related to mitochondrial function and oxidative phosphorylation, including the electron transport chain and ATP synthesis (Supplemental Figure 6D and Supplemental Table 8). Representative genes included *MTCO1*, *SDHD*, *ATP5MC3*, *ATP5MC1*, and *NDUFA12* (Supplemental Figure 5; Supplemental Figure 6E; and Supplemental Table 8), suggesting concurrent shifts in cellular metabolic state.

### Monocyte subpopulation dynamics contribute to longitudinal transcriptional changes

To determine which monocyte populations contribute to these longitudinal transcriptional trends, we examined subpopulation structure at higher resolution. A distinct sub-cluster of 261 CD14+ monocytes, annotated as MARCO+ CCL2+ CLDN5+, was identified (Figure 4A; Supplemental Figure 2E; and Supplemental Table 9). This subpopulation was present in healthy monocytes and jSSc monocytes at baseline and 6 months post-transplant, but was notably reduced in jSSc samples at 12 and 24 months following ASCT (Figure 4B). Upon closer examination, we found that many pathways attenuated over time in all jSSc monocytes (e.g., TNF-α signaling via NF-κB; Figure 4C) were upregulated in this cluster, suggesting that changes in the abundance of specific monocyte subsets may contribute to the observed temporal transcriptional patterns, rather than reflecting uniform shifts across all monocytes.

**Figure 4:**
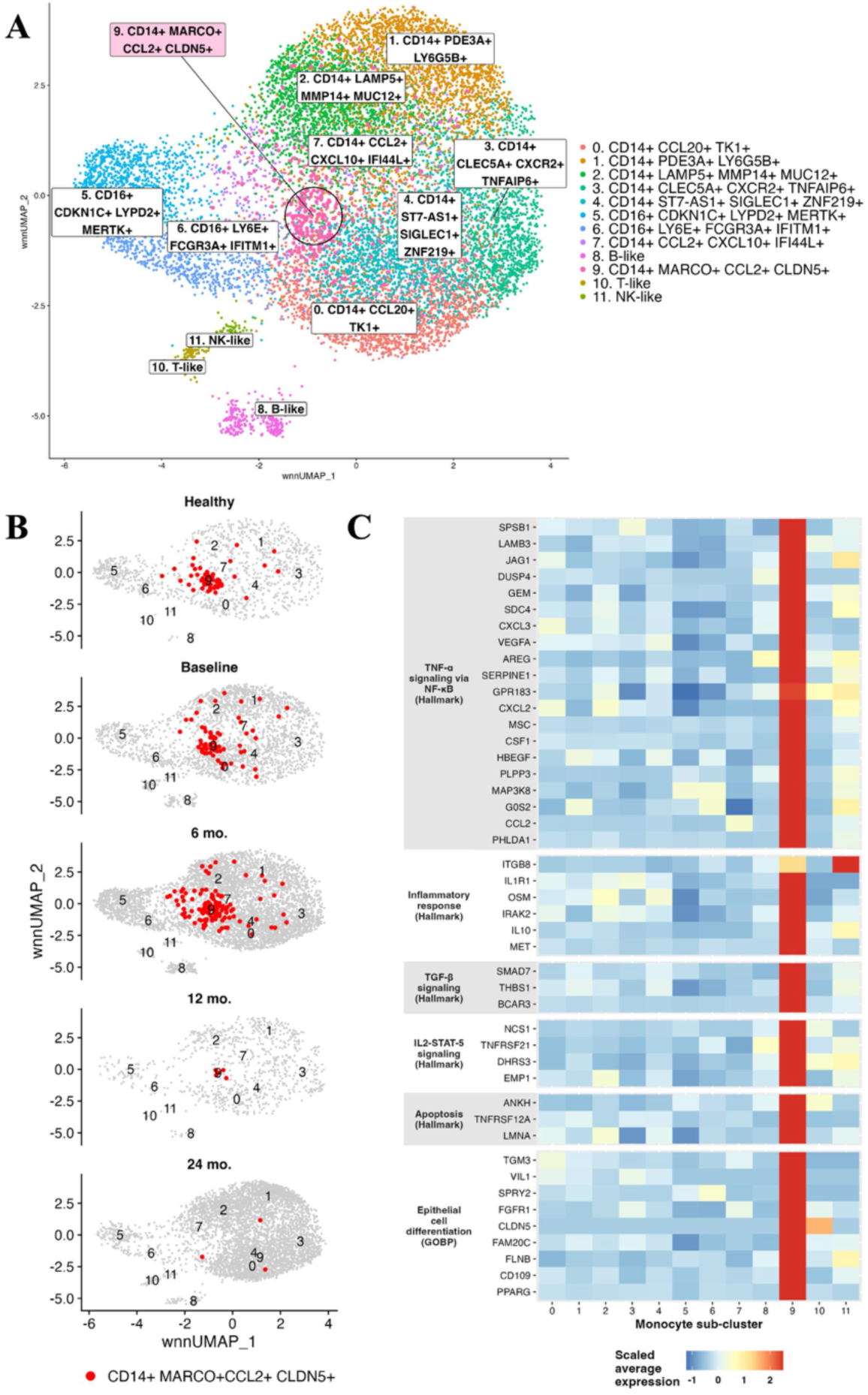
Biological annotation of monocyte sub-clusters and contraction of MARCO+ CCL2+ CLDN5+ population in jSSc monocytes over time since ASCT. A. Biological annotations of jSSc and healthy monocytes. CD14+ MARCO+ CCL2+ CLDN5+ cells (cluster 9) are highlighted as a cluster of interest. B. UMAPs highlighting the CD14+ MARCO+ CCL2+ CLDN5+ monocyte population in healthy controls and jSSc patients over time since ASCT. This population is nearly absent from jSSc patients 12 and 24 months post-ASCT. C. Scaled average expression of key SSc pathway genes in jSSc and healthy monocytes by cluster. These genes are up-regulated in cluster 9.

### Unconventional immune cell populations are identified but require further validation

We identified a small population of cells co-expressing canonical B cell and T cell markers at both the RNA and protein levels (Figure 1 and Supplemental Table 3). These cells passed doublet detection and quality control filters, and were enriched in jSSc samples (95.7% jSSc vs. 4.3% healthy cells, compared to 91.8% jSSc cells vs. 8.2% in the full data set). Similar biphenotypic immune populations have been described in other autoimmune diseases such as multiple sclerosis and rheumatoid arthritis (20). These findings suggest that such cells may represent atypical immune populations in jSSc; however, the possibility of technical artifacts cannot be definitively excluded. We therefore refer to this cluster as T/B biphenotypic cells. RNA and surface protein markers used for annotation are shown in Figure 1, D and E.

Additionally, we identified uncommon monocyte subpopulations, including T-like monocytes, B-like monocytes, and natural killer-like monocytes (Figure 2, A–E; and Supplemental Table 5). These populations comprise very few healthy cells and are seen mainly in jSSc patients at baseline (Figure 2, B and C; and Supplemental Table 6), with a decrease in abundance over time following ASCT.

## Discussion

In this study, we describe longitudinal transcriptional changes in circulating peripheral blood mononuclear cells, with an emphasis on monocytes, in three patients with juvenile systemic sclerosis undergoing autologous stem cell transplantation. Few prior omics studies explicitly investigate the effect of time on the features of interest, but rather perform two-way comparisons between time points or treat time as a categorical variable (21–23). In order to sufficiently capture trends over time, we adapted the offset-pseudobulk model from Lee and Han (16). Using this time-resolved modeling framework, we identified temporal decreases in multiple genes and pathways previously implicated in systemic sclerosis, alongside concurrent shifts in metabolic pathway activity over the two years following treatment.

Several of the most prominent longitudinally decreasing genes, including *SERPINE1* (encoding PAI-1), are well established in the adult systemic sclerosis literature and have been associated with fibrosis, vascular dysfunction, and disease activity. Prior studies have demonstrated increased *SERPINE1* expression in affected tissues (skin and lung) and elevated circulating PAI-1 levels in SSc, supporting its role in disease pathogenesis (17, 18). Similarly, the upregulated TNF-α signaling via NF-κB observed at baseline (pre-ASCT) in jSSc monocytes is consistent with findings from prior studies of SSc serum and keratinocytes (24, 25). In our data set, these inflammatory pathways decreased progressively over time following ASCT, suggesting attenuation of key disease-associated transcriptional programs, which coincided with a dramatic clinical response (26).

Importantly, these transcriptional changes did not appear to be uniform across all monocytes. Subpopulation analyses demonstrated shifts in the abundance of specific monocyte subsets over time, which paralleled the gene expression and pathway-level trends observed in the longitudinal analysis. These findings suggest that changes in cellular composition may contribute to the observed transcriptional dynamics, highlighting the importance of integrating subpopulation structure with gene-level analyses in longitudinal single-cell studies.

In addition to reductions in inflammatory and profibrotic pathways, we observed increased expression of genes associated with oxidative phosphorylation and mitochondrial function over time, which may reflect regeneration and positive biological functions. While not a primary focus of this study, these shifts may reflect broader changes in immune cell metabolic state, potentially consistent with transitions towards less inflammatory or more quiescent phenotypes. Further investigation will be required to better define the biological significance of these metabolic changes in the context of jSSc and ASCT.

We also note that several genes which were upregulated in jSSc monocytes at baseline compared to healthy control monocytes have shown similar patterns in other tissues. For example, *THBS1* overexpression has been observed in the skin of adult SSc patients and is thought to play a critical role in fibrosis (27). Additionally, elevated levels of circulating soluble urokinase plasminogen activator receptor (suPAR), encoded by *PLAUR*, have been observed in adult SSc patients (28).

Clinically, the patients in this cohort demonstrated improvements in multiple organ system domains following ASCT, including skin involvement, gastrointestinal and pulmonary manifestations, and patient-reported outcomes (26). While these clinical improvements occurred in parallel with the observed transcriptional changes, the current study is not designed to establish direct mechanistic links between molecular and clinical responses, and these relationships should be interpreted with caution.

### Limitations

This study has several important limitations. First, the cohort size is small, reflecting both the rarity of jSSc and the limited number of patients undergoing ASCT. Second, although PBMCs were analyzed globally, the present study focused primarily on monocytes, and therefore does not capture longitudinal changes in other immune cell populations that may contribute to disease pathogenesis or treatment response. Third, monocytes comprised a higher-than-expected proportion of the PBMC compartment (48% versus the typical 10–20%), likely reflecting a combination of post-transplant lymphocyte depletion and effects of cryopreservation, which may preferentially impact lymphocyte viability (29–31). These factors should be considered when interpreting cell composition and downstream analyses.

Despite these limitations, this study provides an initial framework for understanding immune dynamics following ASCT in pediatric systemic sclerosis. Our findings highlight coordinated longitudinal transcriptional changes in circulating immune cells and demonstrate the utility of modeling time as a continuous variable in single-cell analyses, especially given multiple post-treatment measurements across therapeutics in autoimmune disease. Future studies in larger cohorts, incorporating additional immune cell populations and independent validation data sets, will be critical to further define these mechanisms and their relationship to clinical outcomes.

## Methods

### Sex as a biological variable

Among the three jSSc patients, there were two females and one male (Supplemental Table 1), while the four healthy controls included three females and one male. These proportions approximate the known female predominance in jSSc, which is about 4:1 (32). Given the limited sample size, sex was not included as a covariate in downstream analyses.

### Study participants and sample collection

All three study participants were between 15 and 21 years of age, had juvenile-onset SSc (onset before the age of 19 years), failed to respond to at least three immunomodulatory therapies, and met ASCT criteria according to severity of skin disease, lung disease, or both (Supplemental Table 1). Participants were enrolled in the National Registry of Childhood Onset Scleroderma (NRCOS; University of Pittsburgh, PRO11060222) and underwent a peripheral blood draw prior to ASCT, and at 6, 12, and 24 months following transplantation. Healthy controls were age- and sex-matched to the patients.

### PBMC CITE-seq

CITE-seq was performed on PBMC samples from all patient time points (baseline, 6, 12, and 24 months post-ASCT) and on matched controls. Approximately 1.5 million cells per sample, with greater than 95% viability, were processed. Cells suspended in Cell Staining Buffer were subjected to Fc receptor blocking using Human TruStain FcX according to the manufacturer’s protocol. All four samples from each subject were stained in a single step with TotalSeq™-C Human Universal Cocktail, V1.0 (BioLegend, Cat No. 399905), along with four individual hashtags (TotalSeq- C0251 anti-human hashtag 1: Cat No. 394661, TotalSeq-C0252 anti-human hashtag 2: Cat No. 394663, TotalSeq-C0253 anti-human hashtag 3: Cat No. 394665, and TotalSeq-C0254 anti-human hashtag 4: Cat No. 394667) to be used for demultiplexing. Following staining, cells were washed three times to remove unbound antibodies and then pooled per subject such that all four time points for a given individual were processed in a single 10x Genomics reaction. cDNA amplification was performed prior to SPRI-based size selection to separate mRNA-derived and antibody-oligo-derived cDNA libraries. Gene expression and antibody-derived tag (ADT) libraries were prepared using 10x Chromium Next GEM Single Cell 5’ Kit v2. VDJ (TCR/BCR) libraries were included as well, but are not analyzed here. Libraries were then sequenced on an Illumina NovaSeq X Plus 25B PE150 platform (Novogene, Sacramento Sequencing Center, CA) and FASTQ files were retrieved from the sequencing provider.

### Sample alignment

Raw FASTQ files were aligned to the human reference genome (GRCh38 v44/Ensembl v110 annotation) using Cell Ranger 8.0.1. In the first step, PBMC libraries were demultiplexed using the *cellranger multi* function, followed by *bamtofastq* (10x). In the second step, alignment for all samples’ RNA and ADT reads was executed with a second pass of *cellranger multi.* The Cell Ranger output was transformed into count matrices in R version 4.4.1 using the Seurat package (version 5.3.0) with the *Read10X()* function.

### Quality control filtering

Multimodal doublet detection was performed on the PBMC CITE-seq samples using the *sccomposite* algorithm (33). Next, ambient RNA correction was applied to all samples with *SoupX* (34). Since this step is meant to address technical variation, it was applied to each of the four PBMC wells, one for each patient and one for healthy controls. Next, standard quality control (QC) filtering was applied. Filtering metrics included number of detected genes, total RNA molecule counts, proportion of mitochondrial RNA, proportion of ribosomal genes, the number of unique antibodies, total ADT counts, and proportion of non-specific binding by isotype control antibodies. For each well, adaptive thresholds were defined as ±3 median absolute deviations (MAD) from the median of each metric (Supplemental Table 2). Following QC, we filtered out genes with low expression, retaining genes with an mRNA count of at least three for at least two different patients at a single shared time point.

### Normalization and scaling

Distinct approaches for normalization and scaling were applied to the RNA and ADT data. For the RNA counts, Seurat’s *NormalizeData() and ScaleData()* functions were applied. The ADT counts were denoised and normalized using the *DSB* algorithm (35). Because the ADT counts did not exhibit a clear bimodal distribution, likely due to challenges with antibody staining, only the first step of the algorithm was applied, which does not rely on distributional assumptions (36).

### Dimensionality reduction and batch integration

For the RNA data, principal component analysis was performed with Seurat’s *RunPCA()* function using the top 2,000 most variable genes as features. Batch effects were corrected using *RunHarmony()* (37), applied separately at the level of individual wells. For the ADT data, Seurat’s anchor-based technique was applied, followed by scaling (*ScaleData()*) and PCA (*RunPCA()*) (38).

### Cell type annotation

A multimodal weighted shared nearest neighbors (WSNN) graph was constructed to integrate RNA and protein data (39). Cells were clustered with the Smart Local Moving algorithm, implemented by Seurat’s *FindClusters()* function and visualized using Uniform Manifold Approximation and Projection (UMAP) via *RunUMAP()*. Cell populations were annotated using canonical RNA and protein markers.

### Longitudinal differential expression analysis in monocytes

After identifying cell types, all 12,744 jSSc monocytes (excluding healthy controls) were extracted and the data were “pseudobulked,” such that total RNA counts for each gene, patient, and time point combination were aggregated. One sample (patient 515 at 12 months) was excluded due to insufficient quality, resulting in 11 evaluable samples out of 12 collected jSSc samples. Aggregated counts were modeled using a generalized linear ‘offset-pseudobulk’ model adapted from Lee and Han, in which the expected gene count was expressed as a function of time since ASCT and patient identity, with an offset for total mRNA count (16),

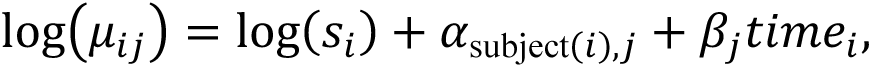

where *μ_i_*_j_ is the expected RNA count for sample *i* and gene *j*; *s_i_* is the total RNA count for sample *i*, summed over all genes; *α*_subject(*i*),j_ is the subject-specific intercept for the subject that sample *i* is from; *β*_j_ is the time coefficient for gene *j*; and *time_i_* is the time in months since ASCT when the sample was collected. Models were fit assuming counts for each gene followed a negative binomial distribution using the R package *glmGamPoi* (40). To identify significant genes, a quasi-likelihood ratio test on *β*_j_ for each gene *j* using the *test_de()* function was applied. After filtering by *P* value (<0.05) and log-fold change (>0.5 or <-0.5), a list of DEGs was obtained in jSSc patients over time following ASCT.

### Pathway enrichment analysis

Differentially expressed genes were stratified by directionality of change over time, distinguishing genes with increasing expression (positive log-fold change) from those with decreasing expression (negative log-fold change) following ASCT. Pathway enrichment analysis was performed on each of the two groups of DEGs using the *enricher()* function from the R package *clusterProfiler* 4.12.6, incorporating gene sets from Hallmark, Gene Ontology (GO) biological process, and GO cellular component databases.

### Statistics

All statistical analyses were performed in R (version 4.4.1). For single-cell analyses, quality control and preprocessing thresholds were applied as described above. Longitudinal gene expression changes in pseudobulked monocyte data were evaluated using a negative binomial generalized linear model with a log-offset for total mRNA count (for a given patient/time point combination across all genes), an indicator variable for patient identity, and time (months post-ASCT) as a continuous predictor. Statistical significance was determined using quasi-likelihood ratio testing. Differential expression required a nominal *P* value < 0.05 and an absolute log-fold change > 0.5. No additional multiple-testing correction was applied given the exploratory nature and small sample size. Pathway enrichment analyses used hypergeometric testing with a significance threshold of *P* < 0.05.

### Study approval

This study was approved by the University of Pittsburgh Institutional Review Board under protocol PRO11060222 (National Registry of Childhood Onset Scleroderma). Written informed consent, and assent when appropriate, was obtained from all participants or their legal guardians prior to inclusion in the study.

## Data availability

Data will be deposited on the Gene Expression Omnibus (GEO) under accession number GSE332819 and will become publicly available upon publication. All analysis code, including R code, Python code, and shell scripts used for the analysis, is available at https://github.com/jkelrod97/jSSc-ASCT-longitudinal-monocytes.

## Author contributions

KST conceived of the study and experimental design and supervised all aspects of the study. AS assisted with experimental design and orchestrated the experiment. JKE and FWT designed the statistical model. JKE carried out all computational processing, with assistance from TH. JKE drafted the manuscript, and KST critically revised the manuscript for important intellectual content. All authors contributed to data interpretation, reviewed the manuscript, and approved the final version.

## Acknowledgments

We thank the patients and families with juvenile systemic sclerosis who participated in the National Registry of Childhood Onset Scleroderma (NRCOS) and generously contributed biospecimens to this study. We acknowledge Kellen Seager and Chaim Sneiderman for critical technical assistance with sample procurement, processing, and experimental execution. We thank Xiangyu Ye for assistance with multimodal cell type annotations. We also thank Haley Havrilla for expert coordination of clinical research activities and patient enrollment.

