## Supplemental Figures for "Longitudinal single-cell modeling reveals monocyte reprogramming in juvenile systemic sclerosis following autologous stem cell transplantation"

**Supplemental Figure 1: Additional UMAPs and marker plots for PBMC annotations**

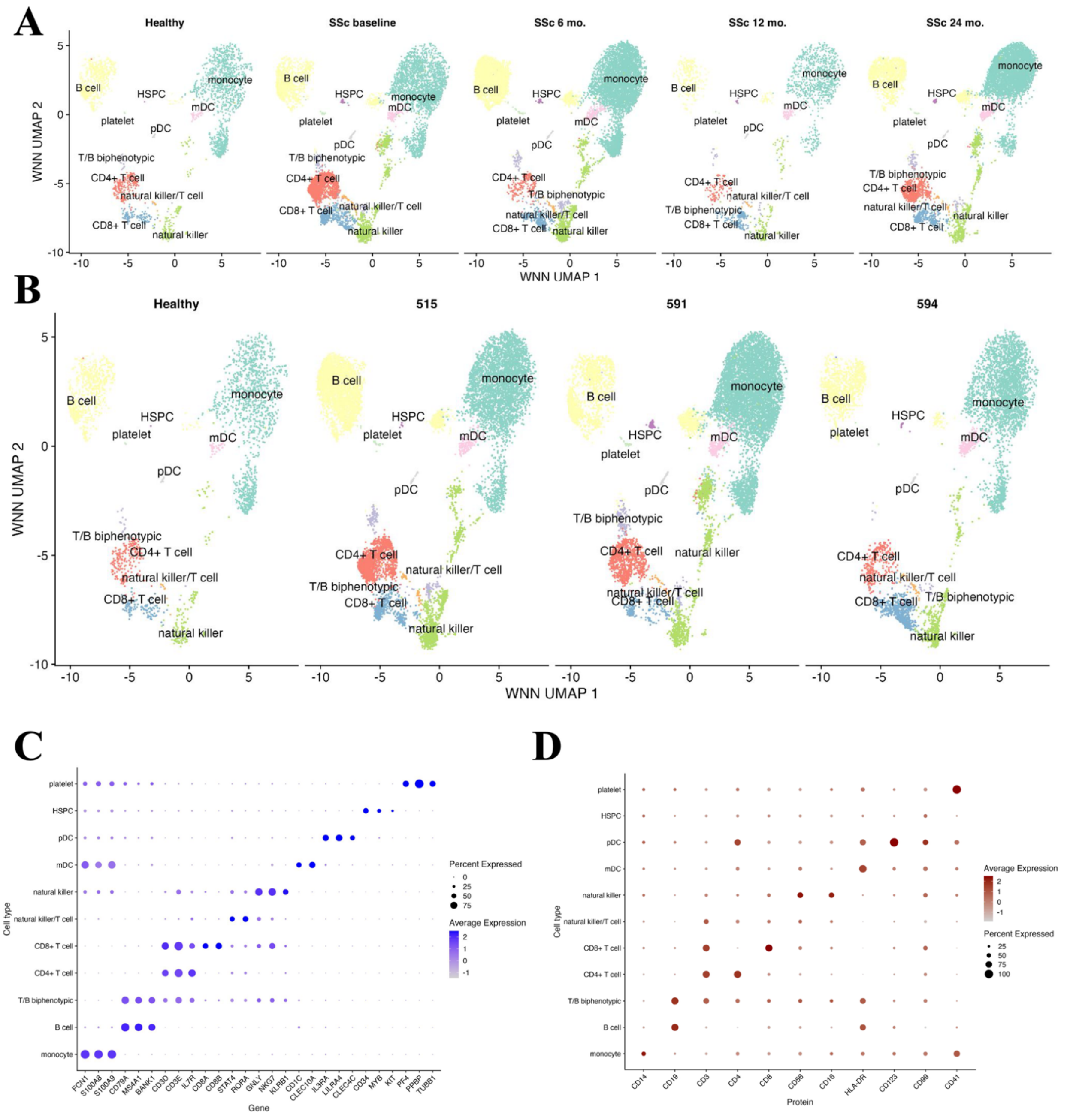


A. UMAPs for all PBMCs, split into healthy controls (left), baseline and time since ASCT for jSSc samples (moving right).
B. UMAPS for all PBMCs, split by library. Healthy combined into one library and individual jSSc subjects' library (combined time points per subject).
C. Dot plot showing RNA markers used for cell type annotation.
D. Dot plot showing ADT markers used for cell type annotation.

**Supplemental Figure 2: Additional UMAPs and marker plots for monocyte annotations**

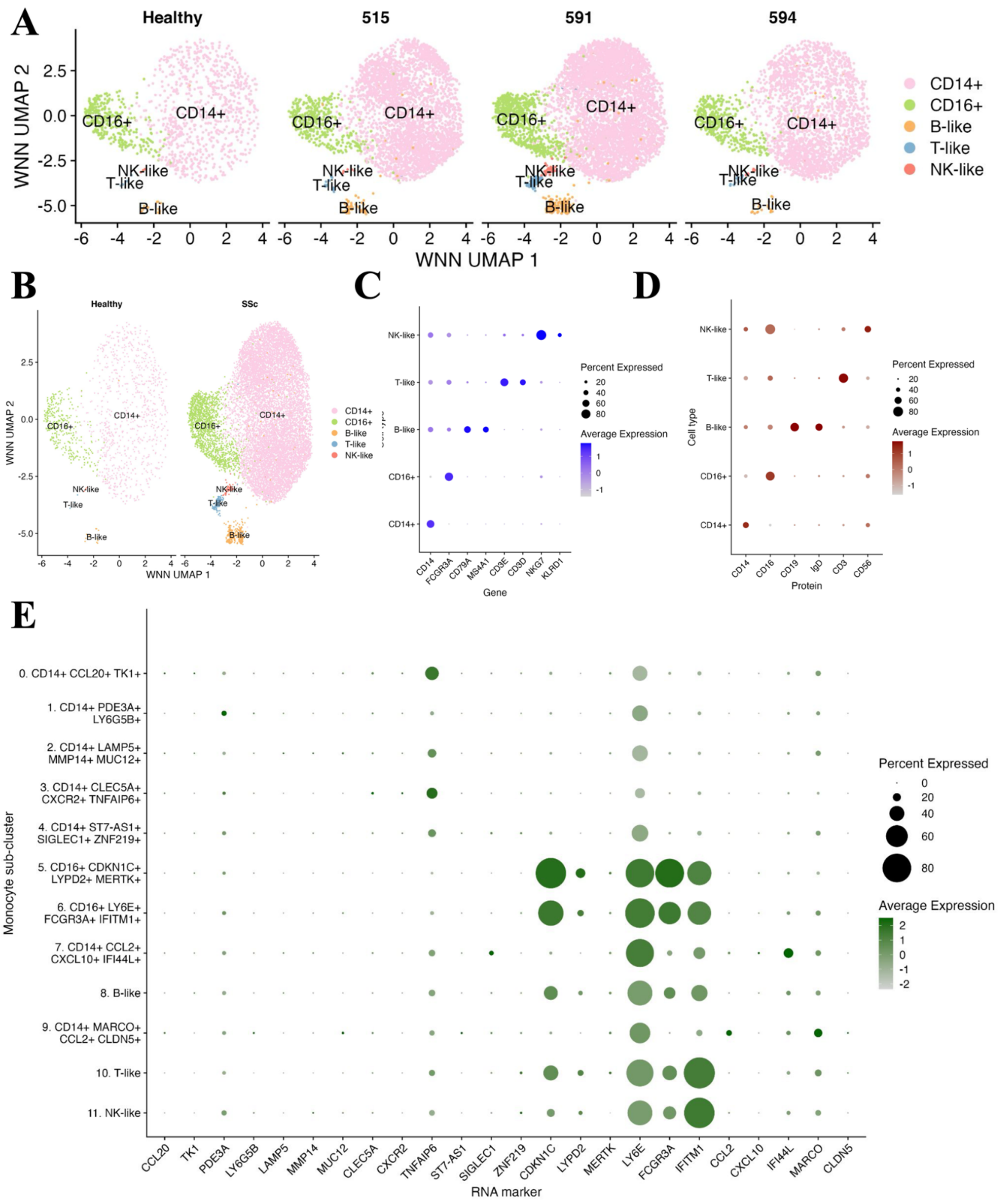


A. UMAPs for all monocytes, split by library.
B. UMAPs for all monocytes, split by health.
C. Dot plot of RNA markers used to identify monocyte subtypes.
D. Dot plot of ADT markers used to identify monocyte subtypes.
E. Dot plot of RNA markers used for biological annotation of monocyte sub-clusters.

**Supplemental Figure 3: Longitudinal gene expression of jSSc monocytes post-ASCT and volcano plots showing DEGs for healthy vs. jSSc monocytes at various timepoints post-ASCT**

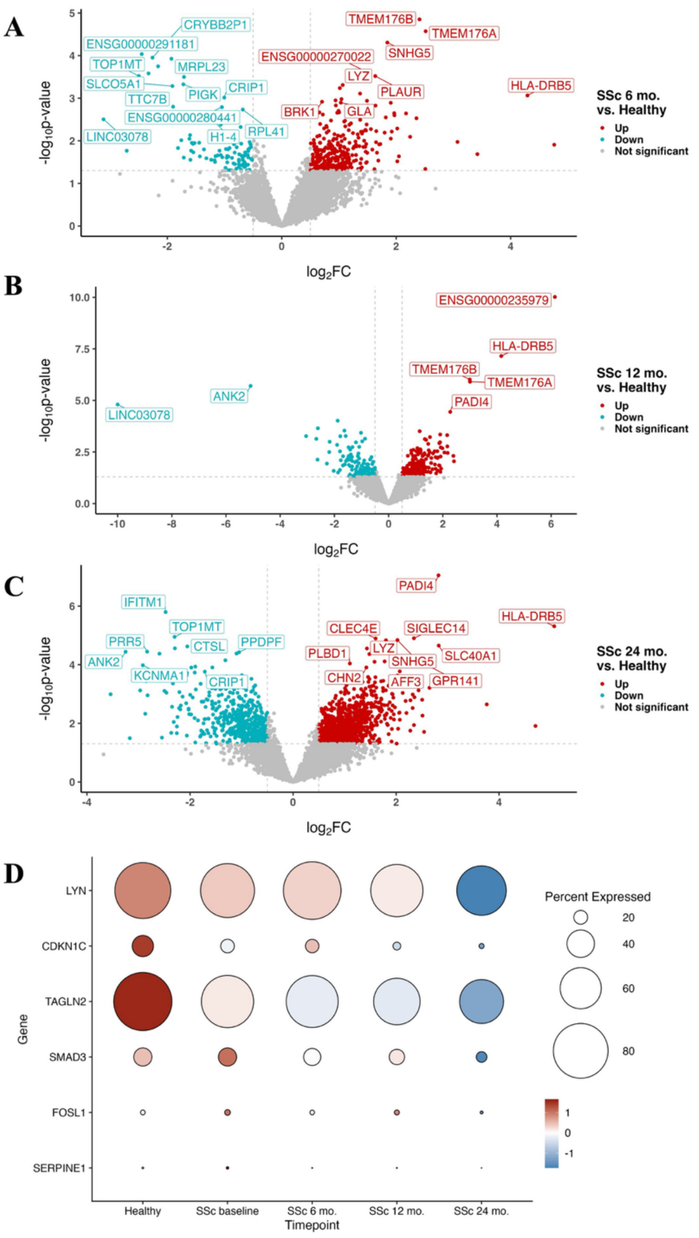


A. Volcano plot showing genes that are up-regulated or down-regulated ($p < 0.05$, $-0.5 < \text{LFC} < 0.5$) in jSSc patients at 6 months vs. healthy controls.
B. Volcano plot showing genes that are up-regulated or down-regulated ($p < 0.05$, $-0.5 < \text{LFC} < 0.5$) in jSSc patients at 12 months vs. healthy controls.
C. Volcano plot showing genes that are up-regulated or down-regulated ($p < 0.05$, $-0.5 < \text{LFC} < 0.5$) in jSSc patients at 24 months vs. healthy controls.
D. Decreasing gene expression patterns in jSSc monocytes over time since ASCT.

**Supplemental Figure 4: Decreasing observed and fitted gene expression in jSSc monocytes over time since ASCT.**

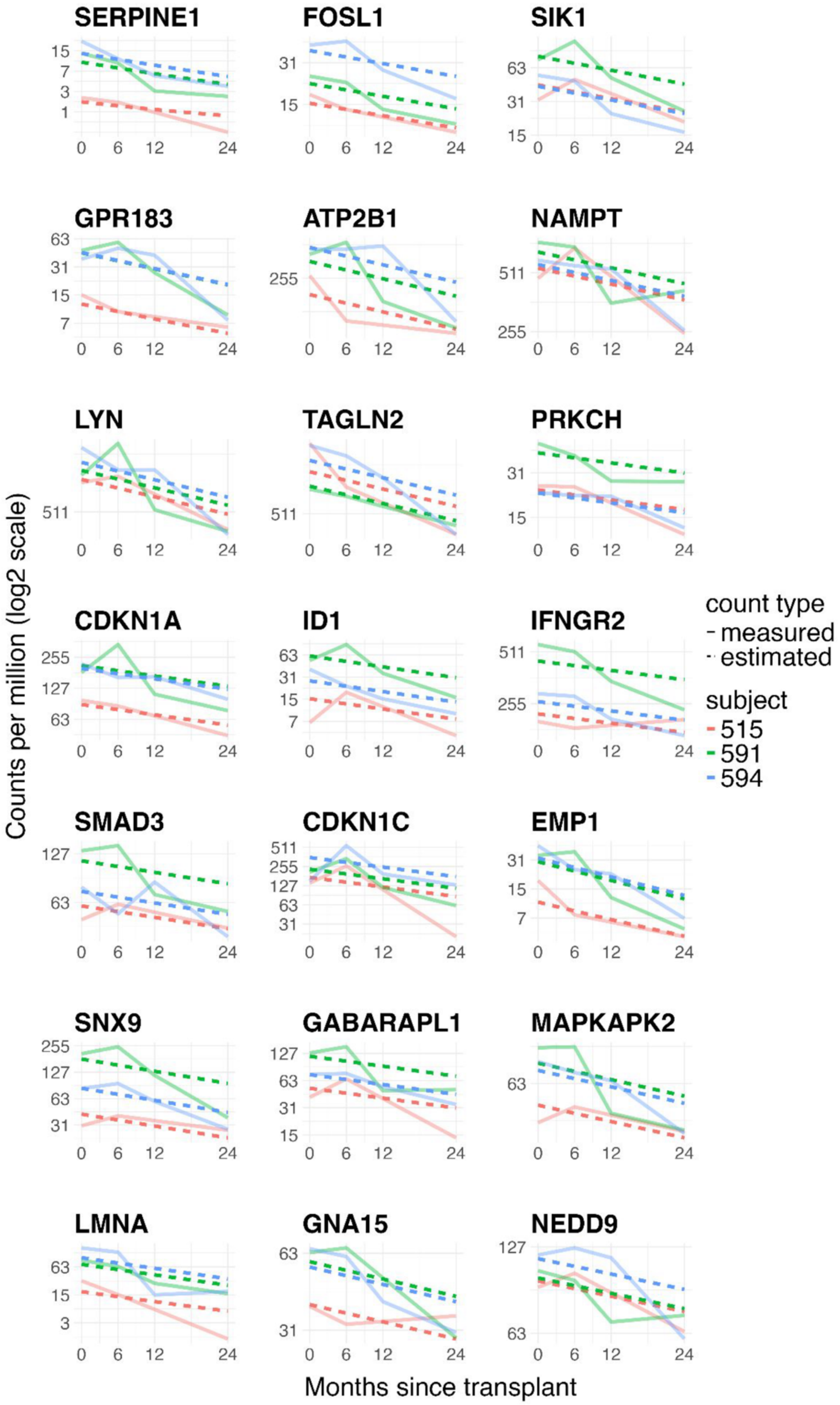

**Supplemental Figure 5: Increasing observed and fitted gene expression in jSSc monocytes over time since ASCT.**

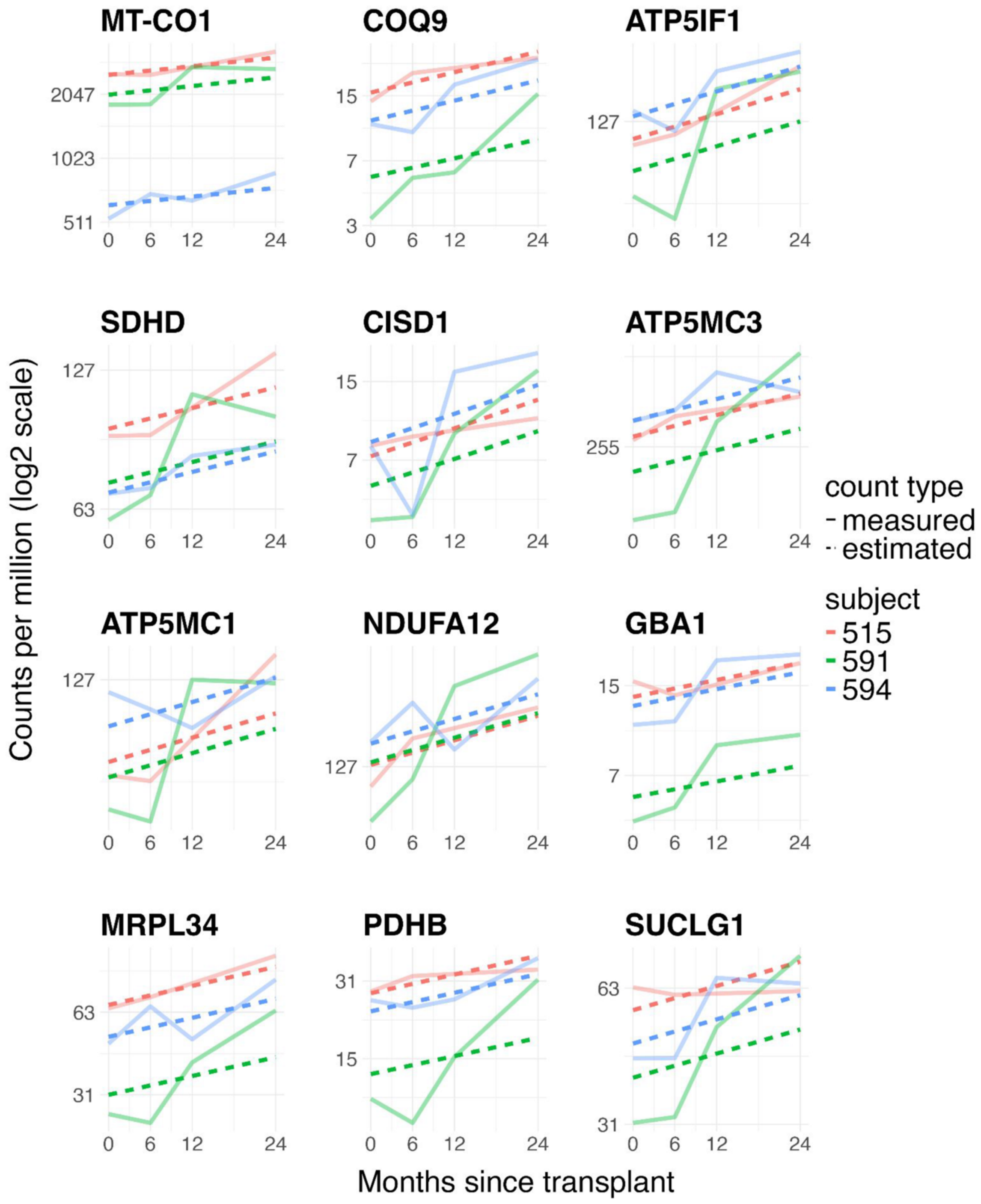

**Supplemental Figure 6: Inflammatory pathway activity in SSc vs healthy monocytes post-ASCT and pathways with increased activity following ASCT**

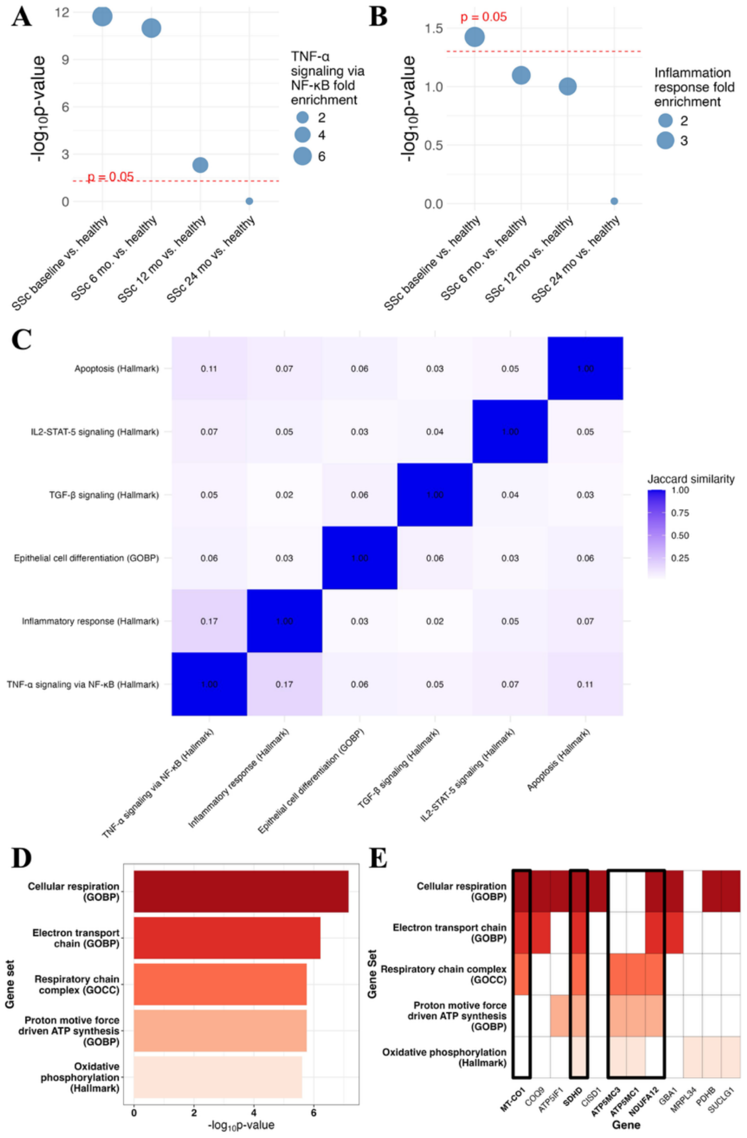


A. Hallmark TNF-α signaling via NFkB pathway activity in SSc monocytes vs. healthy control monocytes over time since ASCT

B. Hallmark Inflammatory response pathway activity in SSc monocytes vs. healthy control monocytes over time since ASCT

C. Jaccard similarity for gene sets with decreased activity in jSSc monocytes over time since ASCT.

D. Gene sets with increased activity in jSSc patients over time since ASCT, selected for biological relevance.

E. Top 5 most significantly increasing genes in each of the pathways in A. Genes appearing in 3+ pathways are highlighted.
